# Determining the defining lengths between mature microRNAs/small interfering RNAs and tinyRNAs

**DOI:** 10.1101/2023.10.27.564437

**Authors:** GeunYoung Sim, Audrey C. Kehling, Mi Seul Park, Cameron Divoky, Huaqun Zhang, Nipun Malhotra, Jackson Secor, Kotaro Nakanishi

**Affiliations:** Center for RNA Biology, The Ohio State University, Columbus, OH 43210, USA; Molecular, Cellular and Developmental Biology, The Ohio State University, Columbus, OH 43210, USA; Department of Chemistry and Biochemistry, The Ohio State University, Columbus, OH 43210, USA; Ohio State Biochemistry Program, The Ohio State University, Columbus, OH 43210, USA

## Abstract

MicroRNAs (miRNAs) and small interfering RNAs (siRNAs) are loaded into Argonaute (AGO) proteins, forming RNA-induced silencing complexes (RISCs). The assembly process establishes the seed, central, 3′ supplementary, and tail regions across the loaded guide, enabling the RISC to recognize and cleave target RNAs. This guide segmentation is caused by anchoring the 3′ end at the AGO PAZ domain, but the minimum guide length required for the conformation remains to be studied because there was no method by which to do so. Using a 3′→5′ exonuclease ISG20, we determined the lengths of AGO-associated miR-20a and let-7a with 3′ ends that no longer reach the PAZ domain. Unexpectedly, miR-20a and let-7a needed different lengths, 19 and 20 nt, respectively, to maintain their RISC conformation. This difference can be explained by the low affinity of the PAZ domain for the adenosine at g19 of let-7a, suggesting that the tail-region sequence slightly alters the minimum guide length. We also present that 17-nt guides are sufficiently short enough to function as tinyRNAs (tyRNAs) whose 3’ ends are not anchored at the PAZ domain. Since tyRNAs do not have the prerequisite anchoring for the standardized guide segmentation, they would recognize targets differently from miRNAs and siRNAs.

## Introduction

More than 2,000 microRNAs (miRNAs) have been reported in humans as of 2019^1^. MiRNAs were defined to be about 22 nt because precursor miRNAs (pre-miRNAs) are processed by Dicer, a molecular ruler which generates size-specific miRNA duplexes^2,3^. The ∼22-nt duplexes are sufficiently long enough to be loaded into Argonaute proteins (AGOs) efficiently. While the passenger strand is ejected, the remaining guide strand and AGO form the mature RNA-induced silencing complex (RISC)^4^. Since both 5′ and 3′ ends of the guide RNA are recognized by the MID and PAZ domains, respectively (Fig. 1a)^5^, the guide acquires the standardized guide segmentation composed of the seed (guide nucleotide positions 2-8, g2-g8), central (g9-g12), 3′-supplementary (g13-g16), and tail (g17-3′ end) regions^6^. The mature RISCs arrange the seed region in the A-form helical structure to enhance the affinity for target RNAs^7^. Guides usually do not involve their central region in target recognition, but their 3’ supplementary region will enhance the affinity for targets when the seed region is mismatched with the target^6^. When RISCs bind to targets through the seed and 3’ supplementary regions, the PAZ domain of the AGO still captures the 3’ end of the loaded guide RNA (Fig. 1a)^8^. However, once the guide-target base pairing proceeds into the tail region, the accumulated topological stress forces the 3’ end of the guide to be released from the PAZ domain^9,10^. As a result, the release of the 3’ end enables the RISC to pair the central region with the target and cleave it if both strands are fully complementary^6,11^. To employ this typical target recognition, the loaded guide must be long enough to maintain the mature RISC conformation. Although the 5′ end of the guide RNA is not accessible to any nucleases due to its thorough recognition between the MID and PIWI domains^5^, the 3′ end of the guide is sometimes released to the solvent and undergoes trimming by 3′→5′ exonucleases or extension by terminal nucleotidyltransferases (Fig. 1a), which generates miRNA isoforms^12,13^. Changing the guide length is known to manipulate the corresponding miRNA’s abundance^14^ and RISC’s half-life, target specificity, and activity^15^.

**Figure 1.**
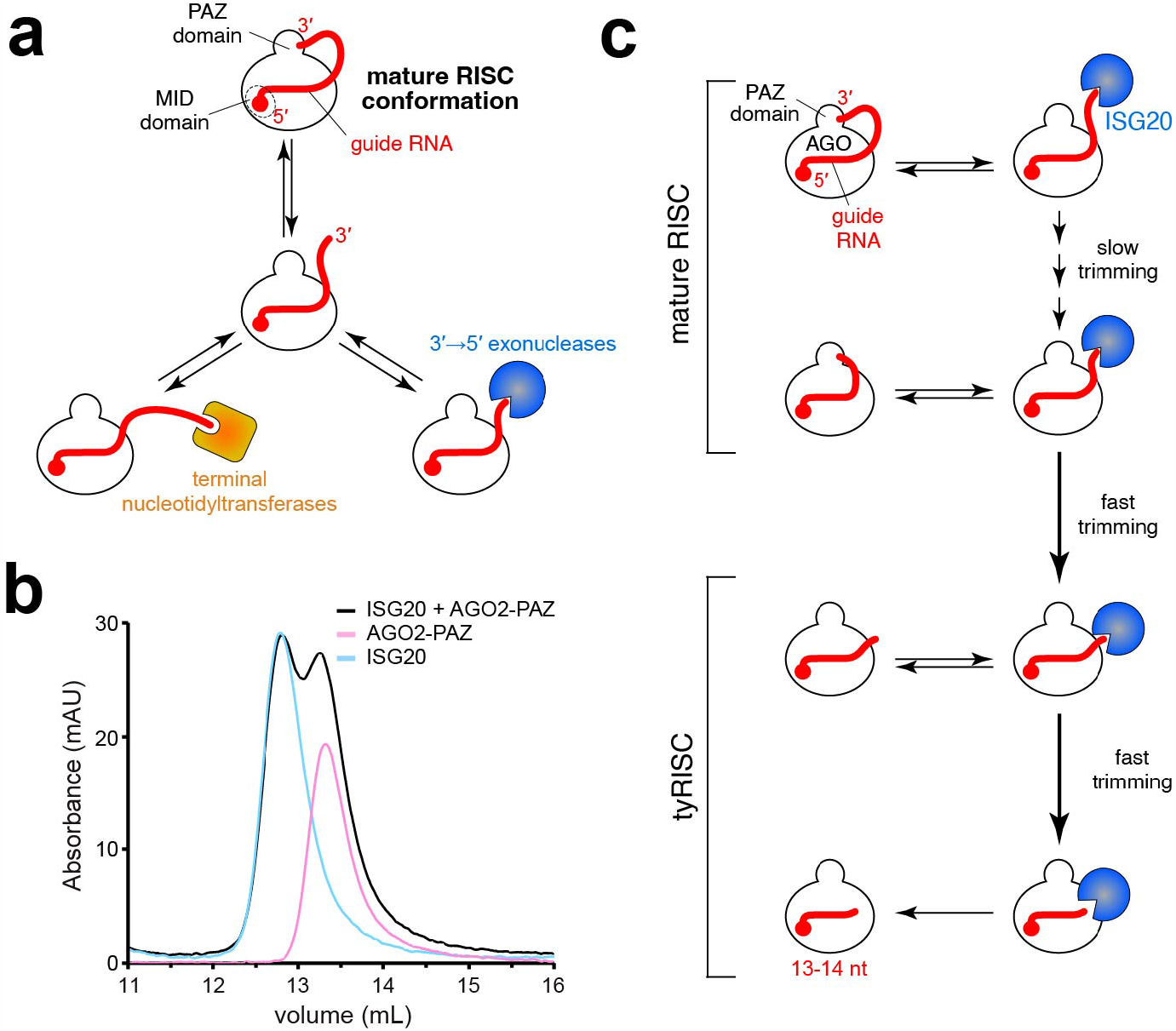
ISG20 is an ideal tool to determine the minimum length of miRNAs. (**a**) Schematic of the competitive interactions of the 3′ end of guide RNA between the AGO PAZ domain and the processing enzymes. In the mature RISC conformation, both 5′ and 3′ ends of the guide RNA are recognized by the MID and PAZ domains. (**b**) Size-exclusion chromatography testing the interaction of ISG20 with the isolated AGO2 PAZ domain. (**c**) Definition of miRNAs and tyRNAs based on the guide length. Mature RISCs become tyRNA-associated RISCs (tyRISCs) upon the conversion of the miRNA to a tyRNA.

Despite its significance, the minimum guide length required for maintaining the mature RISC conformation remains to be studied due to the lack of requisite technology and thus has been a long-standing question.

Meanwhile, AGO-associated guide RNAs shorter than regular miRNAs and small interfering RNAs (siRNAs) have been reported and referred to as tinyRNAs (tyRNAs)^16^. In humans, 17-nt unusually small RNAs were derived from Kaposi sarcoma-associated herpesvirus K12-1 miRNAs^17^, while 15-19-nt tRNA fragments bound to human Argonaute2 (AGO2) to regulate gene expression^18^. In plants, 10-17-nt tyRNAs were discovered inside extracellular vesicles^19^. These previous studies demonstrated that AGO-associated tyRNAs are biologically relevant, but little was known about how these tyRNAs were synthesized^16^. In this context, our recent study discovered a tyRNA-biogenesis pathway in which specific 3′→5′ exonucleases, including interferon-stimulated gene 20 kDa (ISG20), trim 21-23-nt miRNAs down to 13-14-nt tyRNAs^20^. Neither 23-nt miR-20a nor 21-nt let-7a, but rather their 14-nt variants can catalytically activate human Argonaute3 (AGO3)^21^, indicating that tyRNAs play different roles from miRNAs and siRNAs. However, how the size of tyRNAs differs from that of miRNAs and siRNAs has not yet been defined. In this study, we used ISG20 as a tool to determine the defining lengths of tyRNAs and miRNAs/siRNAs.

## Results

### ISG20 competes with the PAZ domain for the 3′ end of guide

Our recent study identified several 3′→5′ exonucleases, including ISG20, capable of trimming AGO-associated miRNAs down to 13-14 nt *in vitro* and *in vivo*^20^. In the current study, we used ISG20 as a tool to determine the minimum guide length required to be captured by the PAZ domain. A previous study reported that small RNA degrading nuclease 1 (SDN1) shortens miRNAs bound to plant AGO^22^. While SDN1 needs to interact with the PAZ domain to initiate the 3′→5′ guide trimming^22^, ISG20 showed no interaction with the isolated PAZ domain of human AGO2 (Figs. 1b and Supplementary Fig. S1). This result demonstrates that ISG20 binds to the free 3′ end only when the PAZ domain transiently releases the 3′ end or cannot capture the 3′ end of short guide RNAs (Fig. 1c). Therefore, we hypothesized that the 3′ end of AGO-associated guides can go back and forth between the PAZ domain and ISG20 (Fig. 1c). The PAZ domain could intermittently protect the 3′ end of the guide from exposure to ISG20 and slow down the trimming rate when the miRNA is long enough to be captured by the PAZ domain. On the other hand, after ISG20 trims the miRNA down to the length of a tyRNA, the 3′ end no longer reaches the PAZ domain and is continuously shortened by ISG20, which accelerates the trimming rate (Fig. 1c).

### miR-20a requires 19 nt to maintain the mature RISC

To validate the potential of ISG20 as a suitable tool, we performed a time-course *in vitro* trimming assay (Fig. 2a and Table 1). After incubation with a 5′-end radiolabeled 23-nt miR-20a, FLAG-tagged AGO2 (FLAG-AGO2) was immobilized on anti-FLAG beads. Any residual free guide was washed out, and ISG20 was added to trim the AGO2-associated guides. The reaction was sampled at different time points, and the trimmed guides were resolved on a denaturing gel. As expected, the trimmed guides were classified into three groups based on their lengths (Fig. 2b). The first group of 19-23-nt guides accumulated well, suggesting that their 3′ end was intermittently captured by the PAZ domain and thus trimmed slowly. The second group of 15-18-nt guides was barely detected, with the 17-nt guides being the least abundant, indicating that their 3′ end was always exposed to ISG20 because the lengths are too short to reach the PAZ domain. The third group of 13-14-nt guides accumulated at the end point of the time course because the nucleic acid-binding channel of RISC sequesters 13-14-nt guides from exposure to ISG20^20^. These results validated our hypothesis and showed that the PAZ domain can capture the 3′ end of the 19-nt miR-20a, but not 18 nt or shorter, if any.

**Table 1.**
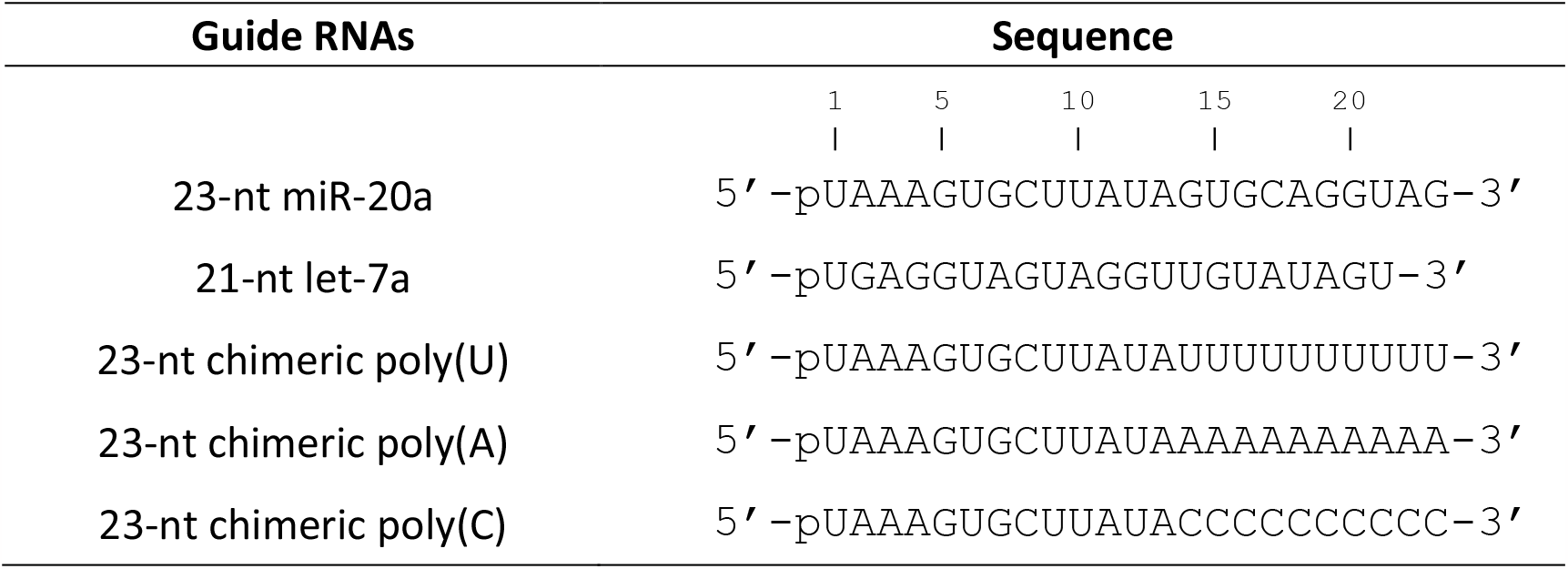
Guide RNAs used in this study.

**Figure 2.**
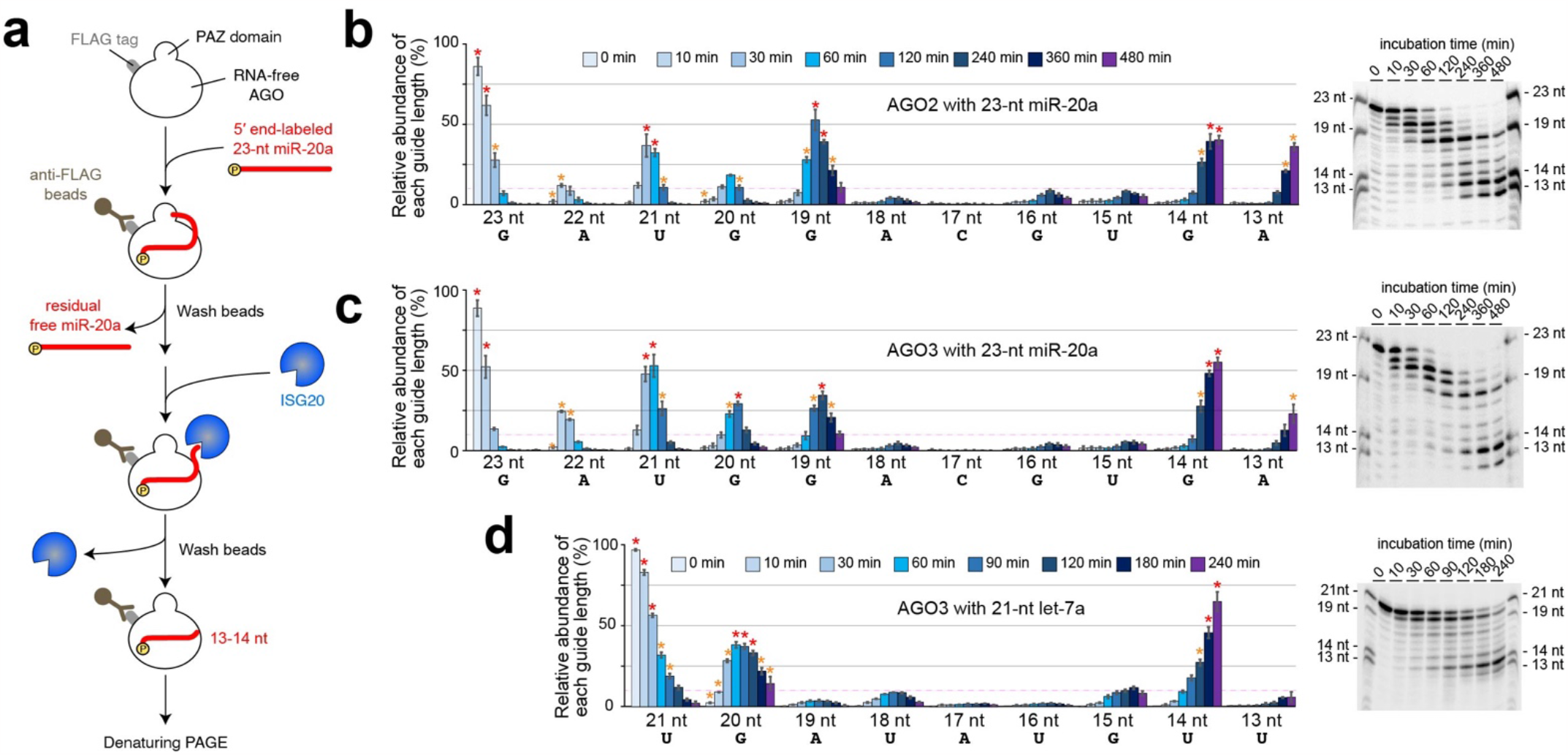
Determining the defining length between miRNAs and tyRNAs. (**a**) Schematic of the *in vitro* trimming assay of FLAG-AGO programmed with a 5′-end radiolabeled 23-nt miR-20a. (**b-c**) Time-course *in vitro* trimming assay of a 5′-end radiolabeled 23-nt miR-20a. *Left*: The relative abundance of 13-23-nt miR-20a bound to FLAG-AGO2 (**b**) and -AGO3 (**c**) over 8 hours was quantified from three independent experiments and shown with mean ± SD. The guide lengths with the highest and second highest abundances at each time point are highlighted with red and orange asterisks, respectively. 10% relative abundance is depicted as dotted pink lines. *Right*: Representative gel images of the guide trimming on FLAG-AGO2 (**b**) and -AGO3 (**c**). (**d**) Time-course *in vitro* trimming assay of 5′-end radiolabeled 21-nt let-7a bound to FLAG-AGO3. *Left*: The relative abundance of 13-21-nt let-7a. *Right*: Representative gel images of the guide trimming.

When compared to the other three AGO paralogs, the AGO2 PAZ domain was reported to have a lower affinity for the 3′ end of the guide because the 3′-end binding pocket possesses Arg315 and His316, instead of lysine and tyrosine, respectively, in the other AGOs^23^. To see whether the trimming pattern varies in paralogous RISCs, the same assay was repeated with FLAG-tagged AGO3 (FLAG-AGO3). While 19-23-nt guides were trimmed slowly, 15-18 nt lengths were barely detected (Fig. 2c). The trimming stopped when the guides became 13-14 nt. The observed trimming profile of FLAG-AGO3-associated 23-nt miR-20a was similar to that of FLAG-AGO2 (Figs. 2b-c). These results demonstrate that ISG20 is a good tool for determining the minimum guide length required to retain the mature RISC conformation and that miR-20a functions as a miRNA as long as the size retains 19 nt.

### let-7a needs 20 nt to stably maintain the mature RISC

The PAZ domain is known to have a preference for UMP, GMP, CMP, and AMP, in this order^24^. Accordingly, we thought that when the 3′ end of the guide has adenine (A), it must be readily released from the PAZ domain and thus trimmed by ISG20 quickly. Such guides would be less prevalent at lengths where A is at their 3′ end, compared with guides of similar lengths that have uridine (U), cytidine (C), or guanidine (G) at their 3′ end. To examine this idea, the *in vitro* guide trimming assay was performed with FLAG-AGO3-associated 21-nt let-7a (Table 1). miR-20a and let-7a have G and A, respectively, at g19, albeit both have G at g20 and U at g21 (Figs. 2c-d). Since the PAZ domain has a higher affinity for GMP than AMP^24^, we surmised that let-7a must not accumulate a length of 19 nt stably, but rather 20 nt. As expected, the 19-nt guide of let-7a remained less than 10% abundance at any time point, unlike miR-20a (Fig. 2d). In contrast, the 18-nt guide of let-7a accumulated slightly more than that of miR-20a (Fig. 2d), which could be explained by the higher affinity of the PAZ domain for UMP over AMP^24^.

However, 20-nt let-7a was always accumulated more than 18-nt at each time point (Fig. 2d), indicating that 20 nt is the minimum length for let-7a to stably maintain the mature RISC conformation. The trimming profile of let-7a also showed the least accumulation at 17 nt (Fig. 2d), which is consistent with the data for miR-20a (Figs. 2b-c). These results suggest that let-7a needs at least 20 nt to employ the standard guide segmentation for target recognition.

### Unbiased guide trimming

The affinity of the PAZ domain for the 3′-end nucleotide impacts the guide trimming rate of ISG20. In addition, the preference of ISG20 for the 3′-end nucleotide of guide RNA would affect the trimming rate as well. We investigated the minimum guide length required for mature RISC in a condition independent of these two biases. To this end, we made three 23-nt chimeric guides whose g1-g13 is derived from that of miR-20a while the g14-g23 is poly(U), poly(A), or poly(C) and named them Chimeric-poly(U), -poly(A), and -poly(C), respectively (Table 1).

Chimeric-poly(G) was highly structured, as reported previously^25^, and could not be used for this assay. Since the g14-g23 of a chimeric guide is composed of identical nucleotides, the 14-23-nt derivatives expose the same 3′-end nucleotide to ISG20 during trimming. Therefore, ISG20 must trim those derivatives at the same rate unless the PAZ domain sequesters their 3′ end. Likewise, the PAZ domain has the same affinity for the 3′-end nucleotides across the 14-23-nt chimeric guide derivatives, so long as it can interact with the 3′ end of the guide. As a result, any observed difference in the accumulation between the 14-23-nt derivatives of a chimeric guide is expected to be attributed to the accessibility of their 3’ end to the PAZ domain.

We programmed FLAG-AGO2 with either of three chimeric guides and tested them for trimming by ISG20. Although the trimming rates were different among the three chimeric guides, which necessitated that a different amount of ISG20 had to be used for each chimeric guide, all of them showed similar trimming patterns in that their 19-23 nt guides accumulated well (Figs. 3a and Supplementary Fig. S2). The 18-nt Chimeric-poly(U) accumulated more than that of Chimeric-poly(A) and -poly(C), as seen in let-7a whose g18 has U. This result further supports that 18-nt guides could stabilize the mature RISC conformation when the g18 is U. Notably, all chimeric guides were the least abundant at lengths of 17 nt, which is consistent with the data for miR-20a (Figs. 2b-c) and let-7a (Fig. 2d). The same trend in guide trimming was observed when FLAG-AGO3 was programmed with these chimeric guides (Figs. 3b and Supplementary Fig. S2). Altogether, our data suggest that the mature RISC conformation remains stable so long as the loaded guide maintains 19 nt or longer.

**Figure 3.**
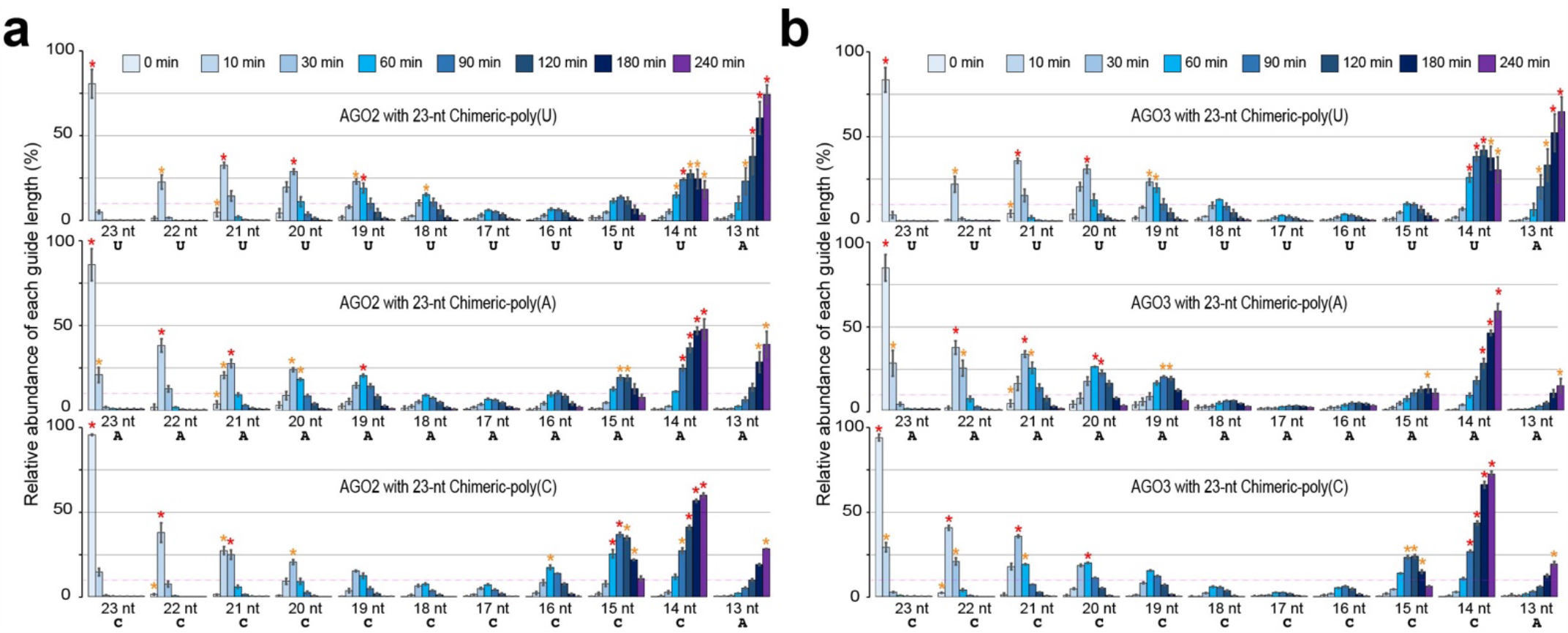
17-nt or shorter guides behave as tyRNAs. (**a-b**) Time-course *in vitro* trimming assay of 5′-end radiolabeled 23-nt chimeric guides on FLAG-AGO2 (**a**) and -AGO3 (**b**). 500, 200, and 80 pmol of ISG20 were used for Chimeric-poly(U) (top), -poly(A) (middle), and -poly(C) (bottom), respectively.

## Discussion

Our unbiased trimming assays indicate that 19-nt or longer guide RNAs maintain the mature RISCs stably (Fig. 4a). This minimum guide length slightly differs depending on the sequence of the tail region due to the preference of the PAZ domain for different 3′-end nucleotides (Figs. 2c-d). In the current study, we defined that miRNAs and siRNAs are long enough to anchor their 3′ end at the PAZ domain. Now we also define guide RNAs whose length is too short to be captured by the PAZ domain as tyRNAs. All of our trimming results showed that 17-nt guides accumulated the least, regardless of guide sequence, demonstrating that 17 nt is sufficiently short enough to keep the 3′ end from being recognized by the PAZ domain. Thus, 17-nt or shorter guide RNAs are defined as tyRNAs, and their ribonucleoprotein complex is termed “tiny RISC (tyRISC)” (Fig. 4b).

**Figure 4.**
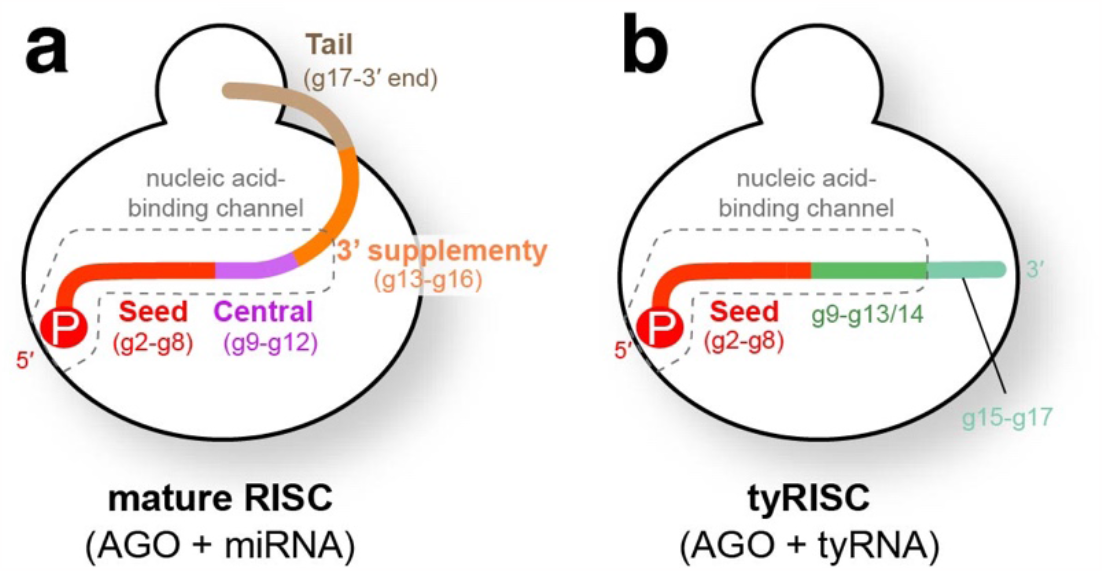
Schematics of mature RISC and tyRISC. (**a**) Mature RISC is composed of an AGO and miRNA. The standardized guide segmentation, the seed (red), central (magenta), 3′ supplementary (orange), and tail (wheat) regions, is shown. The nucleic acid-binding channel is depicted as dotted boxes. (**b**) tyRISC is composed of an AGO and tyRNA. The seed (red) and the g9-g13/g14 (dark green) are protected by the nucleic acid-binding channel from any nucleases. The solvent-exposed g15-g17 (faint green) is easily trimmed by 3′→5′ exonucleases.

Since the nucleic acid-binding channel sequesters the g1-g13/14 from the solvent^20^, the seed of tyRNA is never accessible to any 3′→5′ exonucleases. tyRISCs would use the intact seed region similarly to mature RISCs when recognizing target RNAs. Unlike miRNAs, tyRNAs do not anchor their 3′ end at the PAZ domain of AGO, allowing the g9-g17 to move vigorously.

The g9-g17 of tyRNAs is unlikely to follow the more restrictive standardized guide segmentation of miRNAs. Therefore, after recognizing target RNAs through their seed regions, the tyRISC and mature RISC would treat the target differently. It is known that base-pairing with a fully complementary target occurs sequentially in the order of the seed, 3′ supplementary, tail, and central regions, after which the mature AGO2 RISC cleaves the scissile phosphate group between the t10 and t11^6^. Although AGO3 loaded with specific 14-nt tyRNAs similarly cleaves complementary RNAs between the t10 and t11^21,26^, the actual molecular mechanism must differ from that of mature AGO2 RISCs. Further research is required to understand the mechanisms by which tyRISCs recognize and cleave target RNAs. Future studies should consider this defining size between tyRNAs and miRNAs/siRNAs.

## Methods

### Purification of recombinant proteins

The genes of FLAG-AGO2 and -AGO3 were cloned into pFastBac-HTB (Invitrogen). Their recombinant proteins were overproduced in and purified from insect cells as previously reported and stored in Buffer E (20 mM Tris-HCl pH 7.5, 300 mM NaCl, 0.5 mM TCEP)^26,27^. ISG20 and AGO2 PAZ were cloned into a SUMO-fused pRSFDuet-1 vector (Novagen) and expressed in BL21 Rosetta 2 (DE3) pLysS *Escherichia coli* (*E. coli*) cells (Novagen). All plasmids were verified by Sanger sequencing (OSU Shared Resources). A recombinant protein of ISG20 was purified as previously reported^20^.

*E. coli* cells expressing the isolated AGO2 PAZ domain were lysed with an Emulsiflex C3 homogenizer (Avestin) in Lysis buffer [Buffer A (1x PBS, 500 mM NaCl, 40 mM imidazole, 5% glycerol, 10 mM β-mercaptoethanol) with 1 mM PMSF], followed by centrifugation for 1 hour. The supernatant was loaded onto a 5 mL HisTrap HP column (Cytiva) and washed with Buffer A, followed by elution over a linear gradient to 50% Buffer B (1x PBS, 500 mM NaCl, 1.5 M imidazole, 5% glycerol, 10 mM β-mercaptoethanol). The sample was dialyzed overnight against Dialysis Buffer 1 (1x PBS, 500 mM NaCl, 10 mM β-mercaptoethanol) with Ulp1 protease to cleave the His-SUMO-tag. To remove the tag, the sample was applied to a 5 mL HisTrap HP column (Cytiva) and the flow-through collected for dialysis against Dialysis Buffer 2 (1x PBS, 10 mM β-mercaptoethanol). The sample was loaded to a 5 mL HiTrap Heparin HP Column (Cytiva) and washed with Buffer C (1x PBS, 5% glycerol, 10 mM β-mercaptoethanol) to remove the RNA-bound population. The RNA-free PAZ domain was eluted over a linear gradient to 100% Buffer D (1x PBS, 1 M NaCl, 5% glycerol, 10 mM β-mercaptoethanol) and concentrated by ultrafiltration. The sample was loaded onto a HiLoad 16/600 Superdex 75 pg column (Cytiva) equilibrated in Buffer E (20 mM Tris-HCl pH 7.5, 150 mM NaCl, 5 mM DTT) and peak fractions were concentrated by ultrafiltration. The final concentration was determined by BioSpectrometer (Eppendorf), and samples were flash-frozen in liquid nitrogen to store at – 80°C.

### Binding assay with AGO2 PAZ domain and ISG20

A Superdex 75 increase 10/300 GL column (GE Healthcare) was equilibrated with running buffer (150 mM NaCl, 20 mM Tris-HCl pH 7.5, 2 mM DTT). 100 μmol of the purified AGO2 PAZ domain, ISG20, or a mixture of both was injected to the column. The mixture was incubated on ice for 60 min prior to injection.

### Time-course *in vitro* trimming assay

For the RISC assembly, 20 μM recombinant FLAG-AGO2 or -AGO3 was incubated for 1 hour at 37°C with 2 μM 5′-end radiolabeled guide RNA (23-nt miR-20a, 21-nt let-7a, 23-nt Chimeric-poly(U), 23-nt Chimeric-poly(A), or 23-nt Chimeric-poly(C)) in 1x Trimming Buffer (25 mM HEPES-KOH pH 7.5, 50 mM KCl, 5 mM DTT, 0.2 mM EDTA, and 0.05 mg/mL BSA). 23-nt Chimeric-poly(G), which can form a stable secondary structure^25^, was excluded from the assay. The assembled RISC reaction was pulled down by anti-FLAG M2 affinity gel (Sigma Aldrich), which was pre-washed with 1x PBS twice and 1x Trimming buffer twice, at room temperature for 2 hours with gentle rotation. 50 μL of the 50% slurry of agarose beads were used per 10 μL of reaction volume . Residual free guide RNA was removed by centrifugation, and the coupled beads were washed with immunoprecipitation (IP) wash buffer (300 mM NaCl, 50 mM Tris-HCl pH 7.5, and 0.05% NP-40) 8 times. After removing IP wash buffer by centrifugation, the beads were washed twice with 1x Trimming buffer including 5 mM MnCl_2_. The washed beads were resuspended in 1x Trimming buffer including 5 mM MnCl_2_(25 μL of buffer per 25 μL of packed beads). For 0 min sample, 50 μL of slurry was transferred to a new tube and washed with IP wash buffer 4 times, mixed with 2x urea quenching dye (8 M urea, 1 mM EDTA, 0.05% (w/v) xylene cyanol, 0.05% (w/v) bromophenol blue, 10% (v/v) phenol), and incubated for 1 min at 90°C. ISG20 was added to the rest of the beads and incubated at 37°C for the guide RNA trimming. 500, 200, and 80 pmol of ISG20 per 50 μL of slurry was added to 23-nt Chimeric-poly(U), -poly(A), and -poly(C), respectively.

Because ISG20 trims them at different rates, we used different amounts of ISG20 for each chimeric guide to resolve the trimmed lengths. For each time point, 50 μL of slurry was transferred to a new tube, washed with IP wash buffer 4 times, mixed with 2x urea quenching dye, incubated for 1 min at 90°C, and resolved on an 8 M urea 16% (w/v) polyacrylamide gel. Images were analyzed by the Typhoon Imaging System (GE Healthcare) and quantified by Image Lab (Bio-Rad). All data are presented as the mean ± SD obtained from three replicates.

## Data availability

The sequence information of the plasmids used in the study are available at National Center for Biotechnology Information (NCBI) under the accession ID OR606569, OR606570, OR606571, and OR606572. The data underlying this article will be shared on reasonable request to the corresponding author.

## Acknowledgments

This work was supported by Pelotonia Fellowships (to G.Y.S. and M.S.P.), a Center for RNA Fellowship (to G.Y.S.), and the NIH (R01GM138997 to K.N.).

## Author contributions

K.N. designed the research; G.Y.S., A.C.K., M.S.P., C.D., H.Z., N.M., and J.S. performed experiments and analyzed data. K.N. wrote the paper with input from the other authors. All authors reviewed the manuscript.

